# Combination of *Mycobacterium tuberculosis* RS ratio and CFU improves the ability of murine efficacy experiments to distinguish between drug treatments

**DOI:** 10.1101/2021.12.21.473768

**Authors:** Christian Dide-Agossou, Allison A. Bauman, Michelle E. Ramey, Karen Rossmassler, Reem Al Mubarak, Samantha Pauly, Martin I. Voskuil, Maria Garcia-Cremades, Rada M. Savic, Payam Nahid, Camille M. Moore, Rokeya Tasneen, Eric L. Nuermberger, Gregory T. Robertson, Nicholas D. Walter

## Abstract

Murine tuberculosis drug efficacy studies have historically monitored bacterial burden based on colony forming units of *M. tuberculosis* in lung homogenate. In an alternative approach, a recently described molecular pharmacodynamic marker called the RS ratio quantifies drug effect on a fundamental cellular process: ongoing ribosomal RNA synthesis. Here we evaluated the ability of different pharmacodynamic markers to distinguish between treatments in three BALB/c mouse experiments at two institutions. We confirmed that different pharmacodynamic markers measure distinct biological responses. We found that a combination of pharmacodynamic markers distinguishes between treatments better than any single marker. The combination of the RS ratio with colony forming units showed the greatest ability to recapitulate the rank order of regimen treatment-shortening activity, providing proof of concept that simultaneous assessment of pharmacodynamic markers measuring different properties will enhance insight gained from animal models and accelerate development of new combination regimens. These results suggest potential for a new era in which antimicrobial therapies are evaluated not only on culture-based measures of bacterial burden but also on molecular assays that indicate how drugs impact the physiological state of the pathogen.

## INTRODUCTION

There is an urgent need for shorter treatment regimens for both drug-susceptible and drug-resistant tuberculosis (TB). Murine models have historically been the backbone of preclinical evaluation of TB drugs and treatment regimens (1–3). Pharmacodynamic (PD) monitoring in murine drug experiments conventionally measures colony-forming units (CFU) in lung homogenate. Measurement of 16S rRNA burden has been proposed as an alternative measure of *Mycobacterium tuberculosis* (*Mtb*) burden (4–6). Importantly, neither change in CFU nor rRNA burden during the period that mice are administered treatment accurately indicates the treatment-shortening activity of drugs or regimens (1, 6, 7). Therefore, experiments evaluating the efficacy of multidrug regimens are commonly based on the proportion of mice with microbiologic relapse 12 weeks or more after treatment cessation (8, 9). Because determination of the relapse proportion requires large, resource-intensive mouse experiments sometimes lasting nine to 10 months, the current standard experimental design is a critical bottleneck in TB regimen evaluation. To accelerate regimen evaluation, there is a need for a PD marker or combination of PD markers that indicate the treatment-shortening activity in shorter, less resource-intensive murine experiments without the need for determination of relapse.

We recently proposed a novel molecular PD marker called the RS ratio (10). The RS Ratio measures ongoing ribosomal RNA (rRNA) synthesis in *Mtb* by quantifying the abundance of *Mtb* precursor rRNA (pre-rRNA) relative to stable 23S rRNA. Unlike CFU, 16S rRNA burden and other existing PD markers that enumerate the abundance of *Mtb*, the RS Ratio measures the degree to which drugs and regimens interrupt rRNA synthesis. In the absence of drug treatment, RS Ratio was validated as a surrogate for bacterial replication rate (10). In the presence of drug treatment, RS ratio differentiates individual drug or drug regimen effect in vitro and in vivo and as such, may represent an important new PD marker (10).

In the current work, we used results from three BALB/c mouse experiments to investigate whether three different PD markers (the RS Ratio, CFU, and 16S rRNA burden) provide the same information or measure different biological responses. We asked whether a combination of PD markers measuring different responses distinguishes between treatments better than any single PD marker. Finally, we evaluated the ability of different PD markers and combinations of markers to indicate the treatment-shortening activity of regimens.

## METHODS

We compared three PD markers in three BALB/c mouse experiments in two labs. Experiment 1 evaluated individual drugs to determine whether the RS Ratio, CFU/lung and 16S rRNA burden measure the same or different biological responses. Experiments 2 and 3 evaluated combination regimens to determine whether change in PD markers during the first weeks of treatment distinguishes between regimens and indicates regimen treatment-shortening activity. Efficacy outcomes from Experiments 2 and 3 are reported elsewhere (10, 11).

### Description of BALB/c mouse experiments

Full details of murine protocols are included in Supplemental Information and in other publications (10, 11). Briefly, all three experiments employed female pathogen-free BALB/c mice infected by the same high dose aerosol (HDA) procedure in a Glas-Col inhalation exposure system (12, 13) and treated mice 5 of 7 days a week via oral gavage. Experiments 1 and 3 were conducted at Colorado State University (CSU) using the *Mtb* Erdman strain. Experiment 2 was conducted at Johns Hopkins University using the *Mtb* H37Rv strain.

#### Experiment 1: Individual drug treatments in BALB/c mouse HDA infection model

Treatment began on day 11 and continued for 4 weeks with: bedaquiline (25 mg/kg), ethambutol (100 mg/kg), isoniazid (25 mg/kg), pyrazinamide (150 mg/kg), rifampin (10 mg/kg) or streptomycin (200 mg/kg). Each treatment group had eight mice except for the untreated control which had five mice.

#### Experiments 2 and 3: Multidrug treatments in BALB/c HDA relapsing mouse model

Experiments 2 and 3 used the standard conventional relapsing mouse model described in Supplemental Material (1, 10, 11). In Experiment 2, treatment began on day 14 post-infection with: isoniazid, rifampin, pyrazinamide, ethambutol – (HRZE), rifapentine, moxifloxacin, pyrazinamide – (PMZ), bedaquiline, moxifloxacin, pyrazinamide – (BMZ), or bedaquiline, moxifloxacin, pyrazinamide, rifabutin – (BMZRb). In Experiment 3, treatment began on day 11 with: HRZE using doses identical to Experiment 2, pretomanid, moxifloxacin, pyrazinamide – (PaMZ), bedaquiline, pretomanid, linezolid – (BPaL), or bedaquiline, pretomanid, moxifloxacin, pyrazinamide (BPaMZ). The doses (in mg/kg indicated in subscripts) tested were H_10_, R_10_, Z_150_, E_100_, P_10_, M_100_, B_25_, Rb_10_, Pa_100_, L_100_. The control and treatment regimens each had five mice in Experiment 2, each separate from the mice used for CFU counts in the companion report. The control and treatment regimens each had six mice in Experiment 3.

### Tissue collection

Mice were euthanized the day after the final treatment dose one at a time via CO_2_ asphyxiation. Upper right lung lobes were flash frozen in liquid nitrogen for immediate RNA preservation then homogenized and lysed via beadbeating as described in Supplemental Material. Remining lung lobes were collected for enumeration of CFU.

### Quantification of PD markers

Following RNA extraction and reverse transcription to cDNA, TaqMan qPCR was used to quantify abundance of 16S rRNA as described in Supplemental Material. The RS Ratio was determined in a duplex assay using the QX100 Droplet Digital PCR system (Bio-Rad) as described in Supplemental Material. Primers and probe sequences are in Supplemental Material. CFU burdens were estimated by serial dilutions of lung homogenates and plating on 7H11-OADC agar using 0.4% activated charcoal to prevent drug carry-over as described in Supplemental Material.

### Ethical approval and oversight

Murine experiments were performed in certified animal biosafety level III facilities with appropriate institutional approvals as described in Supplemental Material.

### Statistical analysis

Two-sample Wilcoxon tests were used for pairwise comparisons. For Experiments 2 and 3, a Bayesian sigmoidal E_max_ model was applied using the function “stan_emax” in the rstanemax R package to determine, for individual regimens, the treatment duration that results in 95% cure (T_95_). Then, T_95_ values were compared to establish a rank order of treatment-shortening activity. Lower T_95_ values indicate greater treatment-shortening activity. Description of the sigmoidal E_max_ model is included in Supplemental Material.

Hierarchical clustering was used to evaluate the ability of combinations of PD markers to distinguish drugs and regimens. Differences were considered significant at the 95% confidence level. Analysis was conducted using R (v 3.5.3; R Development Core Team, Vienna, Austria).

## RESULTS

### RS Ratio, CFU, and 16S rRNA burden each measure different biological responses

Treatment with individual drugs affected each of the three PD markers differently (Fig. 1a-c), suggesting that each PD marker measures distinct biological responses. For example, rifampin and isoniazid had similar effects on CFU (*P=0.46*) but rifampin suppressed the RS Ratio 6-fold more than isoniazid (*P*=0.0003). Conversely, isoniazid suppressed 16S rRNA burden 25-fold more than rifampin (*P*=0.0003). Although both CFU and 16S rRNA burden aim to enumerate the quantity of *Mtb*, they did not provide identical information. For example, the effects of isoniazid and bedaquiline on 16S rRNA burden were indistinguishable (*P*=0.3), but bedaquiline reduced CFU 400-fold more than isoniazid (*P*=0.001). Log_10_ decreases and *P-*values for all drugs and all PD markers are included in Supplemental Material Table S1.

**Figure 1.**
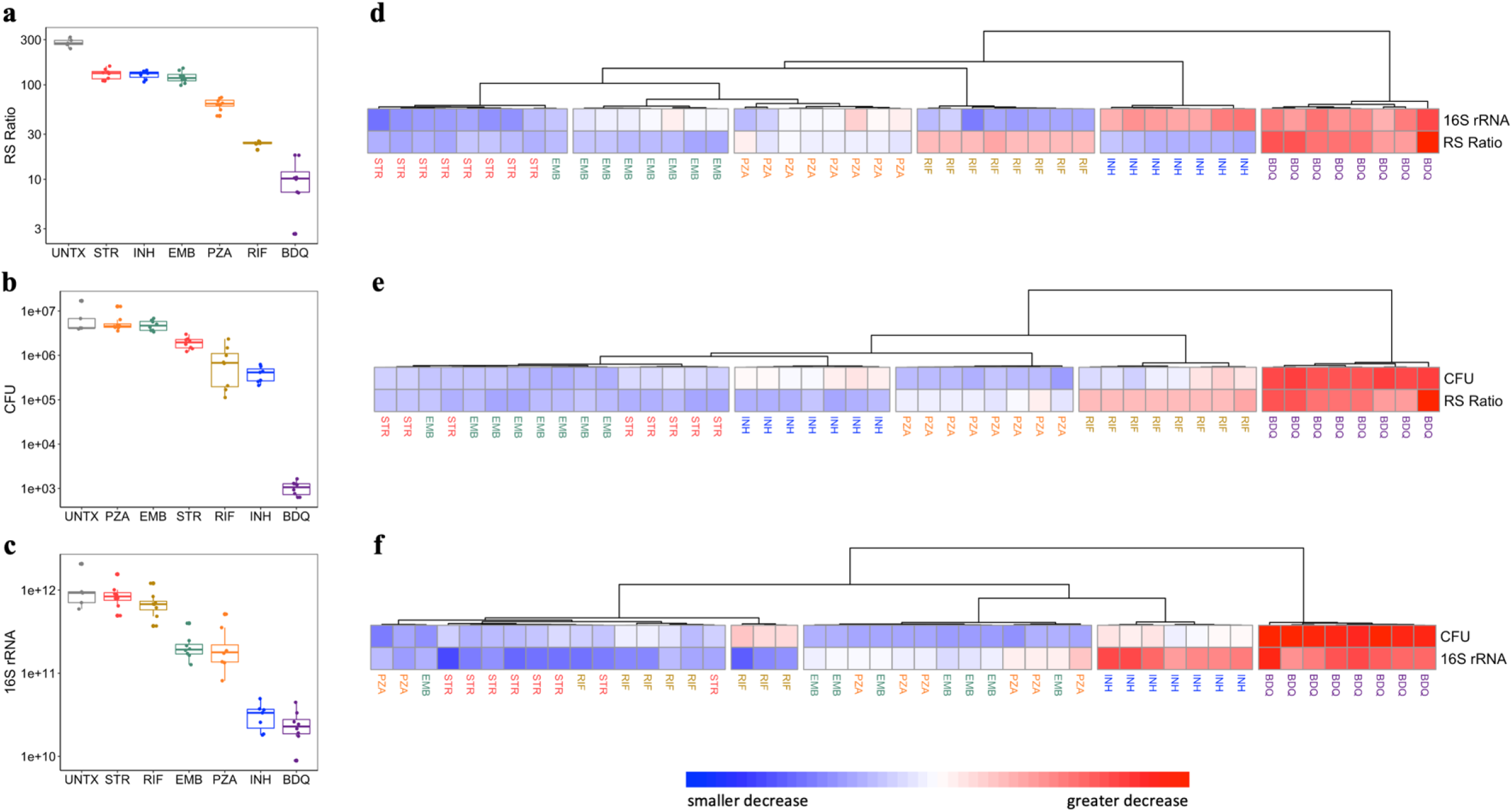
Effect of 4 weeks of treatment 5 of 7 days per week with individual drugs on different PD markers in the BALB/c mouse high-dose aerosol model. Individual drugs had differing effects on (**a**) the RS Ratio, (**b**) CFU and (**c**) 16S rRNA burden. Points and boxes represent untreated control (gray), streptomycin (red), isoniazid (blue), ethambutol (green), pyrazinamide (orange), rifampin (golden) and bedaquiline (purple). Error bars indicate standard deviations. Hierarchical clustering show drug effect on (**d**) 16S rRNA burden and the RS Ratio, (**e**) CFU and the RS Ratio, and (**f**) CFU and 16S rRNA burden in BALB/c mice. Hierarchical clustering was performed using pheatmap in R with the “Ward.D” agglomeration and “Euclidean” distance methods. Rows represent individual PD markers. Columns represent individual mice. Cell values represent log_10_ decrease relative to control. Red, white and blue colors indicate greater, average and smaller decrease, respectively. N=8 mice in each treatment group except for untreated control (N=5). One mouse in INH group was euthanized (Day 18) due to clinical disease resulting in its removal from the analysis.

Each PD marker assessed the rank order of drugs effect differently. For example, isoniazid had the second greatest effect on both CFU and 16S rRNA burden (Fig. 1b-c) but tied with streptomycin for the least effect on the RS Ratio (Fig. 1a). Pyrazinamide tied with ethambutol for the least effect on CFU (Fig. 1b) but had the third greatest effect on 16S rRNA burden (Fig. 1c) and the RS Ratio (Fig. 1a).

### Pairwise combinations of PD markers that include the RS ratio distinguish between drugs better than any individual PD marker

Although no single PD marker was capable of distinguishing between all individual drugs, the distinct effect of drugs could be resolved based on combinations of PD markers that included the RS Ratio. Hierarchical clustering demonstrated that the combination of the RS Ratio and 16S rRNA burden differentiated each drug from every other drug (Fig. 1d). Similarly, the combination of the RS Ratio and CFU differentiated between all drugs with the exception that streptomycin and ethambutol grouped together (Fig. 1e). By contrast, the combination of CFU and 16S rRNA burden largely failed to distinguish between drugs (Fig. 1f). Only isoniazid and bedaquiline were clearly distinguishable; other drugs could not be differentiated based on the combination of CFU and 16S rRNA burden.

### Rank order of treatment-shortening activity in BALB/c relapsing mouse experiments

Experiments 2 and 3 quantified treatment-shortening activity of combination drug regimens in the BALB/c relapsing TB mouse models based on the conventional microbiologic relapse outcome. The relapse outcomes are summarized in Supplemental Material Table S2. Table 1 summarizes T_95_ values. In Experiment 2, the rank order of treatment-shortening activity was: BMZRb (fastest) > BMZ > PMZ > HRZE (slowest) (11). In Experiment 3, the rank order of treatment-shortening activity was: BPaMZ (fastest) > BPaL > PaMZ > HRZE (slowest). The sigmoidal E_max_ model improved model fit and detected significant differences between treatment regimens compared to the hyperbolic E_max_ model (γ = 1) (Supplemental Material, Fig. S1).

**Table 1.** Treatment-shortening activity of diverse regimens in two BALB/c TB relapsing mouse model experiments. Regimen composition, T_95_ and rank order of treatment-shortening activity are shown for Experiments 2 and 3.

| Experiment and regimen | T<sub>95</sub> in weeks<br>[95% confidence interval] | Rank of<br>treatment-shortening activity |
| --- | --- | --- |
| Experiment 2 |  |  |
| bedaquiline, moxifloxacin, pyrazinamide, rifabutin (**BMZRb**) | 6.10 [5.68, 6.31] | 1 |
| bedaquiline, moxifloxacin, pyrazinamide (**BMZ**) | 7.09 [7.05, 7.28] | 2 |
| rifapentine, moxifloxacin, pyrazinamide (**PMZ**) | 8.03 [7.64, 9.82] | 3 |
| isoniazid, rifampin, pyrazinamide, ethambutol (**HRZE**) | 18.63 [18.44, 18.90] | 4 |
| Experiment 3 |  |  |
| bedaquiline, pretomanid, moxifloxacin, pyrazinamide (**BPaMZ**) | 5.59 [5.33, 6.13] | 1 |
| bedaquiline, pretomanid, linezolid (**BPaL**) | 10.00 [9.79, 10.30] | 2 |
| Pretomanid, moxifloxacin, pyrazinamide (**PaMZ**) | 13.12 [12.06, 13.68] | 3 |
| isoniazid, rifampin, pyrazinamide, ethambutol (**HRZE**) | 21.21 [20.70, 21.78] | 4 |

### Correlation of individual PD markers with treatment-shortening rank order

Consistent with Experiment 1, treatment with combination regimens in Experiments 2 and 3 affected the RS Ratio, CFU and 16S rRNA burden differently, confirming that they measure distinct biological responses (Fig. 2a-f). Individually, the three PD markers had variable ability to recapitulate the rank order of treatment-shortening activity of regimens (Fig. 3). After only 4 weeks of treatment in Experiment 2, the RS Ratio by itself matched the rank order of treatment-shortening activity measured many months later (Fig. 3a). CFU by itself did not distinguish the first ranked regimen (BMZRb) from the second ranked regimen (BMZ) (Fig. 3b). 16S rRNA burden by itself did not match the treatment-shortening rank order except that it distinguished between the second (BMZ) and third (PMZ) ranked regimens (Fig. 3c). After 4 weeks of treatment in Experiment 3, the RS Ratio alone did not distinguish between the third (PaMZ) and fourth (HRZE) ranked regimens (Fig. 3d). Likewise, CFU alone did not distinguish between the second (BPaL) and third (PaMZ) ranked regimens (Fig. 3e). Again, 16S rRNA burden alone largely failed to distinguish between treatment regimens (Fig. 3f). Supplemental Material Table S3 summarizes the correlation of individual PD markers with rank order of regimens at the earliest timepoints for both experiments. Supplemental Material Table S4 includes log_10_ decreases for all treatment regimens and timepoints.

**Figure 2.**
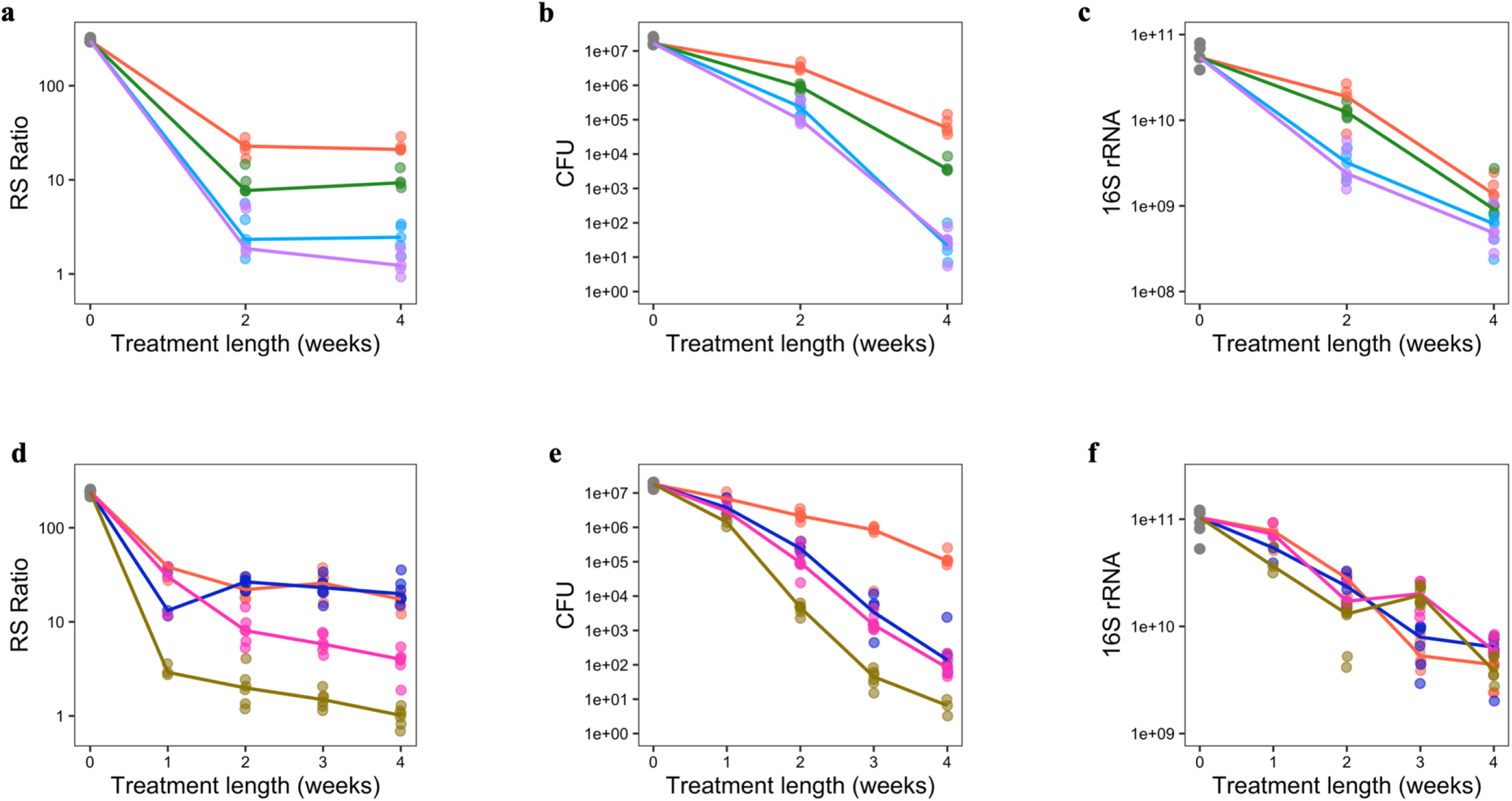
Differing effects of treatment on three PD markers during the first 4 weeks of treatment using data from two BALB/c TB relapsing mouse model experiments. RS Ratio, CFU and 16S rRNA burden were measured in lung homogenate of untreated mice (grey) and mice treated with BMZRb (purple), BMZ (light blue), PMZ (green) and HRZE (orange) in Experiment 2 (**a, b, c**), and with BPaMZ (golden), BPaL (pink), PaMZ (blue) and HRZE in Experiment 3 (**d, e, f**). Dots represent values from individual mice. Solid lines connect median values. All graphics use a log_10_ scale for the Y axis. The control and treatment regimens each had 5 mice (Experiment 2) and 6 mice (Experiment 3).

**Figure 3.**
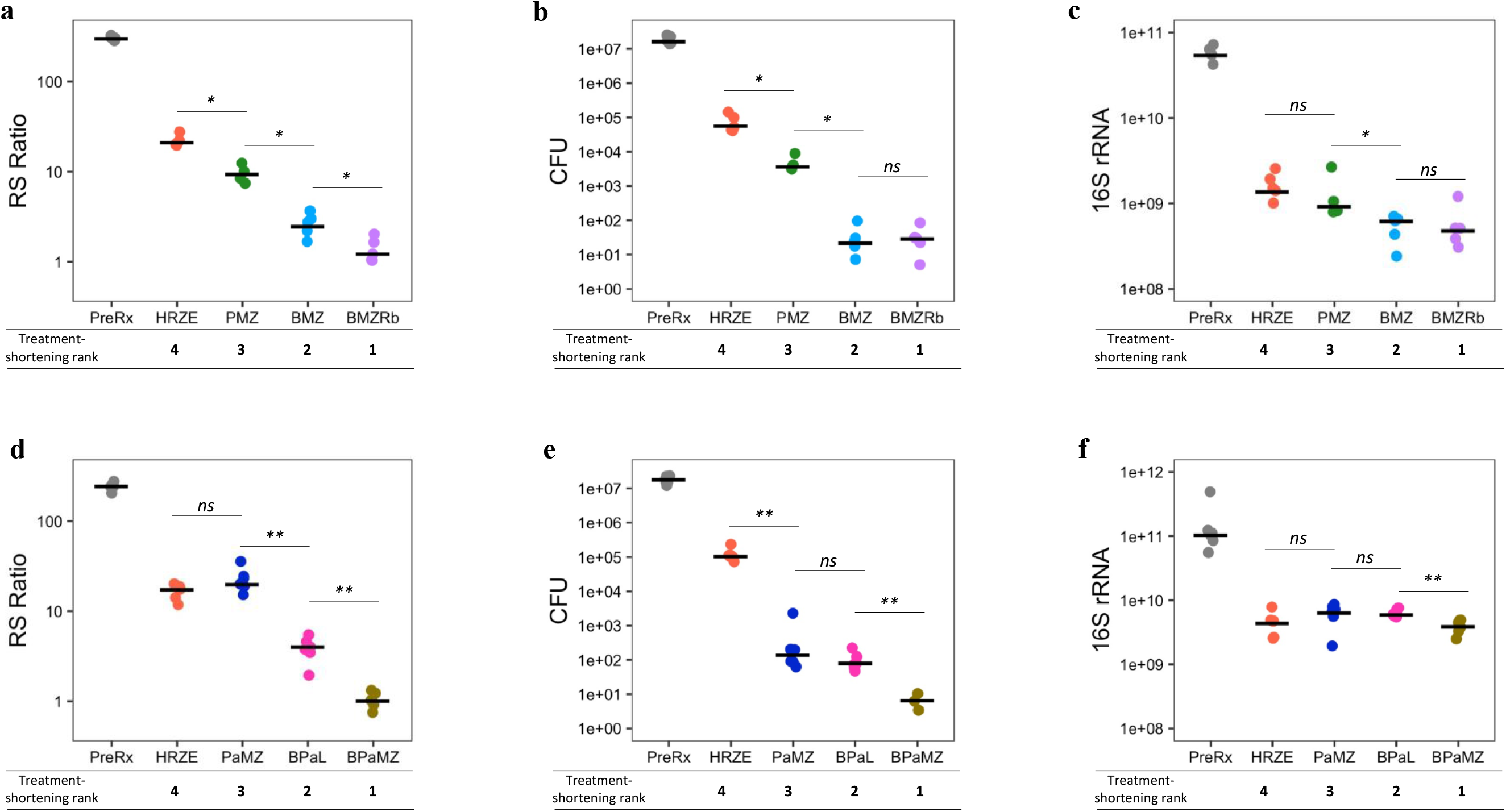
Correlation of RS Ratio, CFU and 16S rRNA burden with treatment-shortening rank order after 4 weeks of treatment using data from two BALB/c TB relapsing mouse model experiments. RS Ratio, CFU and 16S rRNA burden were measured in lung homogenate of untreated mice (grey) and mice treated with BMZRb (purple), BMZ (light blue), PMZ (green) and HRZE (orange) in Experiment 2 (**a, b, c**), and with BPaMZ (golden), BPaL (pink), PaMZ (blue) and HRZE in Experiment 3 (**d, e, f**). Dots represent values from individual mice. Bars represent median values. *P*-value symbols are as follows: ns is non-significant, * is *P*-value <0.05, ** is *P*-value <0.01. All graphics use a log_10_ scale for the Y axis.

### A combination of the RS Ratio and CFU improves distinction and classification of treatment regimens

Combining different types of PD markers assisted in distinguishing the distinct effects of different regimens. After 4 weeks of treatment, the combination of the RS Ratio with CFU near-perfectly distinguished between regimens in Experiment 2 (Fig. 4a) and perfectly distinguished between regimens in Experiment 3 (Fig. 4d). The degree to which regimens decreased CFU and RS Ratio appeared concordant with treatment-shortening activity (Fig 4a, 4d). By contrast, the combination of 16S rRNA burden with either the RS Ratio (Fig. 4b, Fig. 4e) or CFU (Fig. 4c, Fig. 4f) failed to distinguish between treatment regimens.

**Figure 4.**
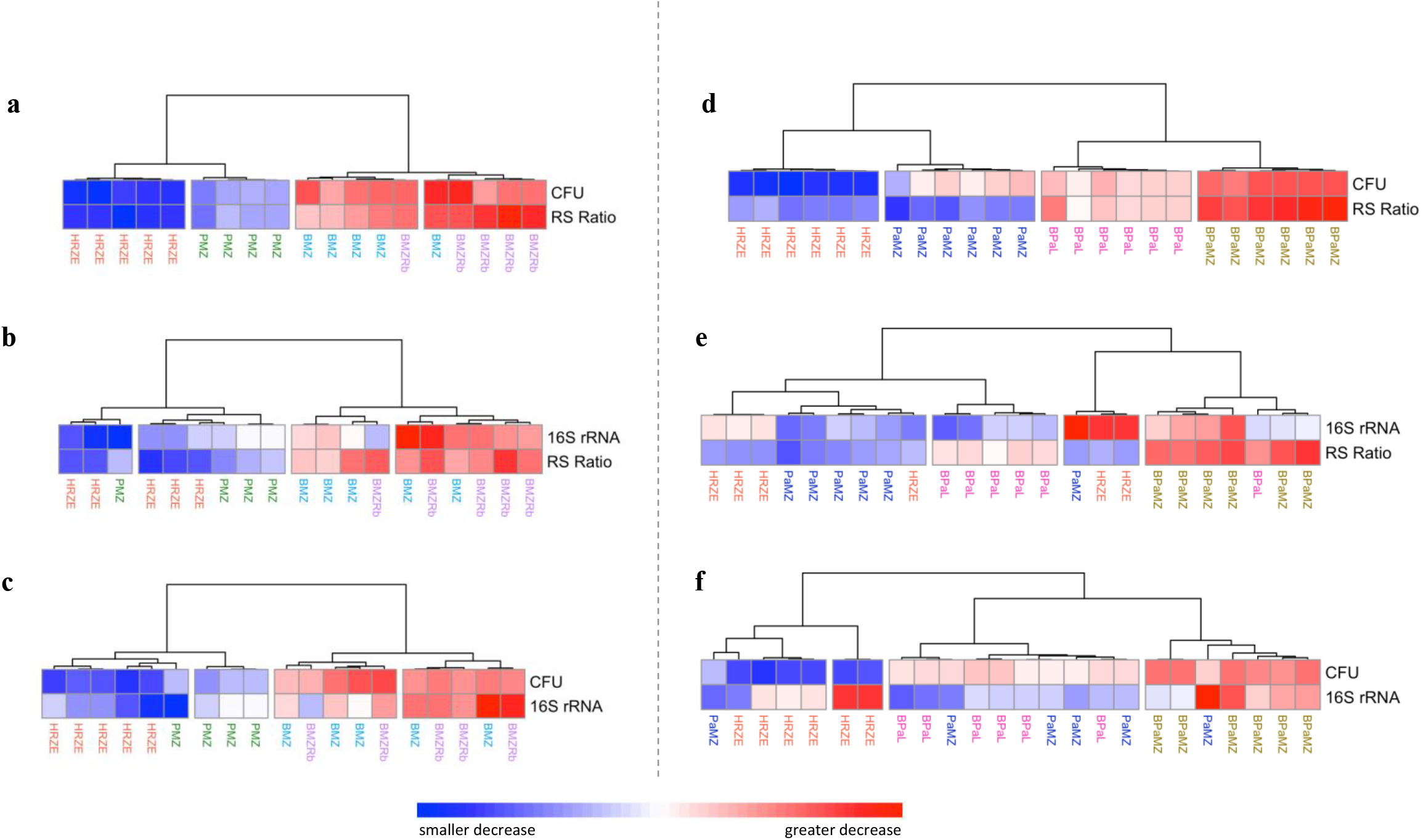
Distinguishing different regimens based on three PD markers measured after 4 weeks of treatment using data from two BALB/c TB relapsing mouse model experiments. Combination of PD markers are shown for Experiment 2 (**a, b, c**) and Experiment 3 (**d, e, f**). Hierarchical clustering was performed using pheatmap in R with the “Ward.D” agglomeration and “Euclidean” distance methods. Rows represent individual PD markers. Columns represent individual mice. Cell values represent log_10_ decrease relative to control. Red, white and blue colors indicate greater, average and smaller decrease, respectively. The control and treatment regimens each had 5 mice (Experiment 2) and 6 mice (Experiment 3). RS Ratio could not be quantified in one mouse treated with PMZ in Experiment 2 resulting in its removal from the analysis.

## DISCUSSION

Our analysis of three BALB/c mouse experiments, conducted at two different institutions using different infecting strains, demonstrated that the RS Ratio, CFU, and 16S rRNA burden each measure different biological responses to drug treatment. The RS Ratio is a non-culture-based assay that provides orthogonal information and correlates with regimen treatment shortening activity. Combining different PD markers enhanced distinction between treatments, relative to any single marker alone. The combination of the RS Ratio with CFU showed the greatest ability to recapitulate the rank order of regimens, providing proof of concept that assessment of regimen treatment-shortening activity within the first weeks of treatment may be possible. Development of an early accurate measure of treatment-shortening activity has potential to transform the design of murine efficacy studies, thereby accelerating evaluation of new more potent regimens.

CFU has been a standard historical marker in murine studies despite widely recognized limitations. Our results reinforce previous evidence that change in lung CFU during the first weeks of treatment in mice does not indicate the treatment-shortening activity (1). Perhaps relatedly, McCune and colleagues in the 1950s (14, 15) and more contemporary investigators (16–21) have shown that CFU quantifies only the subset of the *Mtb* population that is capable of growth on solid media and often does not detect the last remaining viable bacilli that determine the treatment duration necessary to prevent relapse in mice. These limitations highlight the potential value of gathering alternative information from murine drug studies and motivated our development of molecular measures of treatment effect.

rRNA has been proposed as a means of enumerating the entire *Mtb* population, including subpopulations capable and incapable of colony formation on solid media (6). de Knegt *et. al*. previously described a striking divergence between reduction in CFU and reduction in the Molecular Bacterial Load Assay (MBLA, a measure of rRNA burden) in BALB/c mice. For example, de Knegt found that, after 8-12 weeks of treatment with HRZE, CFU decreased >100-fold more than MBLA. Our current experiments demonstrated a similar disconnect in which CFU decreased more than 16S rRNA burden. It remains unclear whether the sustained high rRNA burden during treatment indicates the presence of a continuing large non-culturable *Mtb* population or detection of residual rRNA from dead *Mtb*. Our experiments confirmed de Knegt’s observation that change in rRNA burden largely fails to distinguish between regimens with different treatment-shortening potency in mice.

Unlike CFU and rRNA which estimate bacterial burden, the RS Ratio was designed to measure an alternative property: the degree to which drugs and regimens interrupt rRNA synthesis. Each of our three experiments demonstrated that the RS Ratio provides information that is orthogonal to CFU or rRNA burden. Experiment 2 showed that the RS Ratio was able to measure the effect of adding single drug (Rb) to a potent combination (BMZ), a difference that was not identifiable based on CFU. Change in the RS Ratio correlated with the treatment-shortening activity of regimens. These observations suggest that the RS Ratio may be a valuable new non-culture-based tool for preclinical efficacy evaluation.

This study also provides proof of concept that different readouts of drug effect (*i.e*., CFU and RS Ratio) can be complementary and their combination may be more informative than either PD marker alone. A next step will be development of a composite outcome incorporating CFU and the RS Ratio to improve early efficacy assessment in mice. This would require a development phase in which both CFU and the RS Ratio are collected in additional relapsing mouse trials testing diverse regimens. These results would enable parameterization of a composite outcome and evaluation of the composite CFU-RS Ratio (quantified during the first treatment weeks of treatment) as a surrogate for subsequent relapse. If prediction of relapse is validated, a composite CFU-RS Ratio assay would enable higher throughput murine screening studies in which a large number of regimens is tested in one-month studies to “funnel down” to top candidates that can then proceed to traditional, lengthy, resource intensive, relapsing TB mouse model experiments. Availability of a method that reliably predicts treatment-shortening efficacy based on responses during the first weeks of treatment would alleviate a key bottleneck in the preclinical TB drug evaluation process.

This study has several limitations. First, an inherent challenge to evaluating the predictive value of PD markers in mice is that mice can only be sacrificed once. Because it is not possible to measure PD markers early in treatment and the relapse outcome in the same individual mouse, predictive modeling is inherently limited. Second, as noted above, this report demonstrates proof of concept based on two relapsing mouse studies, establishing a starting point. Parameterization and validation of a composite CFU-RS Ratio will require additional relapse studies with more diverse regimens.

In summary, this analysis highlights the potential to harness multiple different types of PD markers to extract greater insight from animal models and accelerate development of new combination regimens. New molecular tools like the RS Ratio offer potential for a new era in which antimicrobial therapies are evaluated not only on culture-based measures of bacterial burden but also on molecular assays that indicate how drugs impact the physiological state of the pathogen.

## DATA AVAILABILTY

All primary data is included in the Supplemental Material.

